# Effectiveness of a network Open House model to recruit trainees to post-baccalaureate STEM programs

**DOI:** 10.1101/2024.01.08.574670

**Authors:** Scott Takeo Aoki, Lindsay Lewellyn, Sarah Justice, Sarah Mordan-McCombs, Neetu Tewari, Jorge Cantu, Robert Seiser, Ahmed Lakhani, Jennifer R. Kowalski

**Author notes:** **Consent Statement:** Pre- and post-Open House surveys were approved by the Butler University Institutional Review Board (IRB) on September 18, 2023. Mentor and MSI surveys were approved by the Butler University IRB on December 19, 2024.

## Abstract

Post-baccalaureate (post-bac) programs can be instrumental in strengthening science training and expanding STEM career opportunities for junior trainees. Many of these sponsored programs are designed to increase research exposure for underrepresented minorities, including African American, Hispanic, Native American, and first-generation college students, among others. Recruiting trainees to post-bac programs can be challenging for reasons including a lack of awareness about available programs. To address this gap, an Open House event was created with the goal of raising awareness and generating interest among potential students for future post-bac programs. Students were recruited from partnering Minority Serving Institutions (MSIs) to attend a two-day event at a Primarily Undergraduate Institution (PUI) and a Research-Intensive (R1) institution. During the visit, students toured both campuses, learned about various post-bac programs and research opportunities, and interacted with faculty, current graduate students, and a former post-bac scholar. Transportation, lodging, and meals were provided. Participants completed voluntary pre- and post-surveys. Results indicated that attendees, the majority of whom were underrepresented minorities in STEM, left with a stronger understanding of post-bac programs and how these experiences could support their future careers in STEM and that students’ attendance at the event made it more likely they would apply to available post-bac programs. Mentor and MSI faculty survey responses highlighted their strong support for participating in future recruitment events. These findings demonstrate that in-person Open House events, built on collaborative partnerships across institutions, are an effective strategy for increasing awareness and encouraging participation in post-bac training programs— particularly among underrepresented student populations.

## Introduction

Starting a career in science depends on extensive hands-on experience. For many, laboratory research experience begins in their high school or undergraduate education, but for others, obligations outside the classroom prevent them from experiencing bench research firsthand. This challenge is often observed with students who identify as underrepresented minorities in science or who have come through a community college system (1, 2), and it can limit individuals belonging to these groups from obtaining the lab research experience necessary for graduate programs or employment in science, technology, engineering, and mathematics (STEM) careers. For example, graduate schools look for meaningful research experience in their candidates. In many programs, matriculating graduate students are years past their undergraduate education (3), giving them time to obtain relevant research experience that they might not have had the opportunity to pursue while working towards their bachelor’s degree. Developing opportunities for students to gain experience after their undergraduate training is central to recruiting a diverse, balanced population to the STEM workforce, but many of those who would benefit most from these opportunities may be unaware of their existence or benefits underscoring the need for targeted outreach and accessible pathways into research training.

Post-baccalaureate (post-bac) programs are one to two-year funded, research-intensive training experiences designed to prepare trainees for graduate school and STEM careers. For example, the National Institutes of Health (NIH) Postbaccalaureate Research Education Program (PREP) program was established in 2000 to support post-bac trainees at a variety of research institutions across the country (4). This program evolved new strategies to promote readiness for STEM graduate school (5, 6) and has been incredibly successful. Of recent PREP scholars, 65-97% enter graduate school programs, and Ph.D. completion rates are > 65% above the rates reported for underrepresented groups in the life sciences (6-8). The American Cancer Society (ACS) and National Science Foundation (NSF) also developed post-bac programs with similar structural models (9, 10) reflecting a critical role these program play in supporting research training for individuals historically underrepresented in science. While a funded research experience outside of schooling promises more opportunity to recruit a breadth of students from a wide demographic, but a persistent challenge faced by post-bac programs is how to reach trainees who may not be familiar with the benefits of these programs or who do not have access to established pathways leading to a successful STEM career.

An Open House event invites candidate trainees on site to introduce a program and present opportunities available to them. Although traditionally associated with K-12 or undergraduate recruitment, these events are inherently flexible and can be effectively adapted to support recruitment at the later stages of the educational pathway. by design and can be impactful well past the traditional K-12 use of such events. Targeted, personal Open House-like events can be helpful in recruiting individuals from specific demographics, like those who identify as female and African Americans (11). Students considering various undergraduate programs also have identified Open House events as an effective recruiting tool (12). Universities note that Open House events are a chance to present a positive image to visitors (13). Open Houses are a chance for real human connection, which can showcase the advantages of an educational program to groups of people missed through other advertising campaigns.

In this study, an Open House event was developed to introduce the benefits of post-bac programs, with an emphasis on reaching students from groups underrepresented in the biological sciences (14) with little previous research experience. Faculty and students from Research-Intensive Institutions (R1s), Primarily Undergraduate Institutions (PUIs), and Minority-Serving Institutions (MSIs) who form collaborative research networks are effective in undergraduate biology training (15), and personalized referrals are among the most effective strategies for recruiting students from underrepresented minority groups to STEM graduate school (11). In consideration of these factors, an event was crafted that leveraged the strengths of faculty partnerships across a network of MSI, PUI, and R1 institutions. The effort created an experience that reached a cohort of students from underrepresented minority groups in science and presented post-bac programs as a viable steppingstone for a STEM career.

## Materials and Methods

### Open House Event and Student Survey Formats

Recruitment for the Open House was performed through advertising and word of mouth. The advertising flyers, which were tailored to be institution-specific, were created in Canva (Canva, Sydney, Australia; www.canva.com) and contained a QR code linked to a Google Form (Google; Mountain View, CA; www.google.com) for registration. Students were selected on a first come, first served basis. Partnering MSIs were given first access to registration, followed by students at the hosting institutions. While 30 students could have been accommodated, 17 students were recruited to the event, with 15 attending on both days. Students and faculty from their home institutions were responsible for arranging travel to Indianapolis, IN. Hotels were reserved through Butler University, the primary hosting institution. Butler University’s Provost’s Office funded all events. These funds covered the hotel costs, food, pens, programs, folders for attendees, and mileage reimbursements. MSIs covered the cost of transportation rental to bring their cohorts, as needed.

#### Day 1

Students and faculty arrived at Butler University, a PUI in Indianapolis, IN. Prior to scheduled events (**Fig S1**), students completed an anonymous pre-survey (**Supplemental Information 1**), approved by a Butler University IRB (*Approval date: Sept. 18, 2023*) and administered by Qualtrics (Qualtrics; Provo, UT), taking approximately 10-15 minutes to complete. This survey requested information regarding the participant’s demographics, science experiences, and familiarity with and interest in post-bac programs. Seventeen students initiated the pre-survey, but only 15 students completed it. Respondents were not required to complete all the questions; thus, there is some variation in the number of responses per question. Following survey completion, students then learned of the opportunities for post-bacs and those with science graduate degrees (e.g., M.S., Ph.D.), research opportunities at local PUIs, and resources available at Butler University. A tour of the Butler University campus was made available for those interested. Visiting students and faculty then were taken to dinner with faculty interested in hosting post-bacs and with graduate students from the Indiana University School of Medicine (IUSM), an R1 institution. Visiting faculty and students stayed at a local hotel sponsored by the program.

#### Day 2

Students and faculty visited Indiana University School of Medicine; Indianapolis, IN (**Fig S1**). They were given an overview of an established post-bac program (https://iprep.iupui.edu/index.html) and research at Indiana University and interacted with a graduate student panel assembled by the local chapter of the Society for the Advancement of Chicanos/Hispanics and Native Americans in Science (SACNAS). Tours of the Centers of Electron Microscopy and Proteome Analysis facilities were given. A sponsored lunch was provided with Indiana University School of Medicine faculty members and graduate students. Visiting students were prompted to complete a Qualtrics exit survey consisting of the similar questions regarding post-bac programs (**Supplemental Information 2**). A total of 13 students completed this exit survey, approved by the Butler University IRB (*Approval date: Sept. 18, 2023*). As with the pre-survey, respondents were not required to answer all questions, again leading to some variation in response numbers per question.

### Mentor/Co-Mentor and MSI Faculty Post-Open House Surveys

An anonymized survey was distributed to prospective faculty mentors from Butler University, DePauw University, Marian University, IUSM, and science faculty at partner MSIs. The survey assessed their motivations for and interest in supporting post-baccalaureate training programs. Faculty from PUIs and R1 universities were asked about their experience mentoring students from historically underrepresented backgrounds. MSI faculty were asked about how post-bac training programs aligned with their professional goals, the challenges of balancing recruitment efforts with other responsibilities, and the potential impact of these programs on their students. The surveys were approved by the Butler University IRB (*Approval date: December 19, 2024*) and administered via Qualtrics (Provo, UT). Completion time was approximately 5–10 minutes.

### Data Analysis

Anonymized pre- and post-event student survey data were aggregated separately and analyzed for statistical significance in GraphPad Prism version 10.4.2 for MacOS (GraphPad Software, Boston, Massachusetts USA). Figure 1A and B data were analyzed using a two-tailed Mann Whitney U test to compare pre- and post-survey Likert score means converted to a 1-5 scale. Figure 1C data were analyzed using a Wilcoxon signed-rank test with the neutral response (3.0 = *neither agree nor disagree*) at the middle of the 1-5 Likert scale set as the theoretical median value. Figures 2 and 3 data were converted to percentages of respondents and graphed. Figures were made using Prism and Adobe® Illustrator® (Adobe, San Jose, CA). Qualtrics data for all survey questions are included in **Supplemental Information 1-4**.

**Figure 1.**
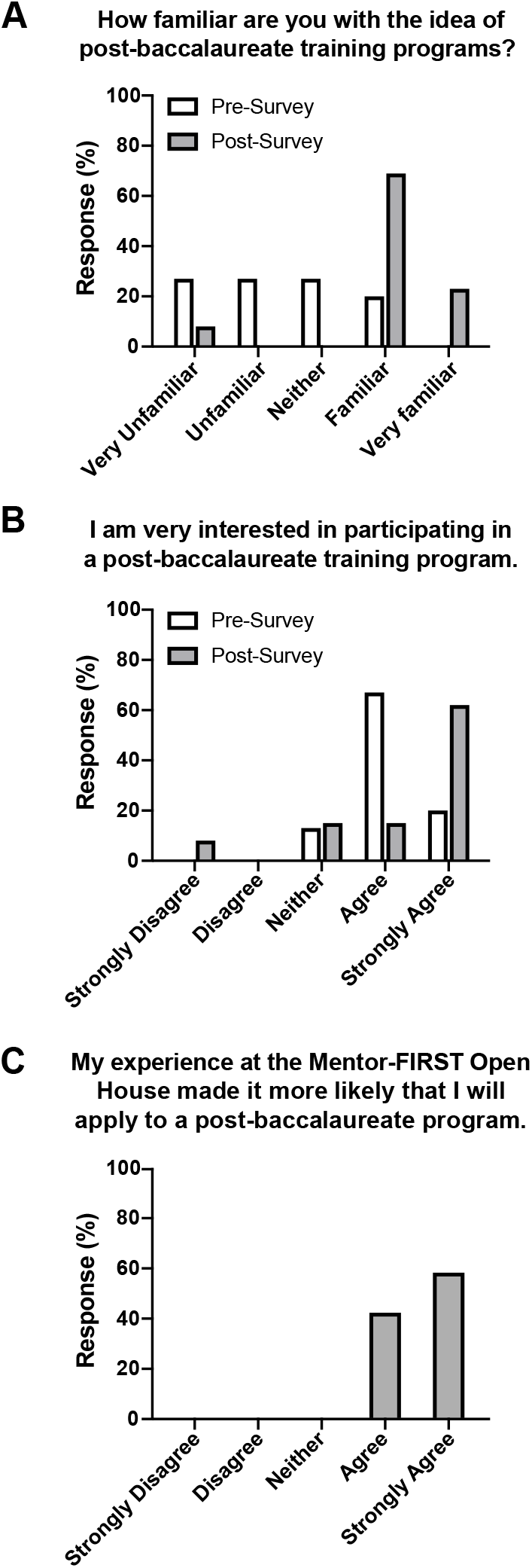
Effectiveness of an Open House event in educating and promoting post-baccalaureate programs. **(A, B)** Pre- and post-event survey responses regarding (A) student familiarity with post-baccalaureate training programs and (B) student interest in participating in a post-baccalaureate training program (N = 15 pre; N = 13 post). **(C)** Post-survey responses regarding the impact of the Open House event on the likelihood of their future application to a post-baccalaureate training program (N = 12). See *Results* text for statistical analysis.

**Figure 2.**
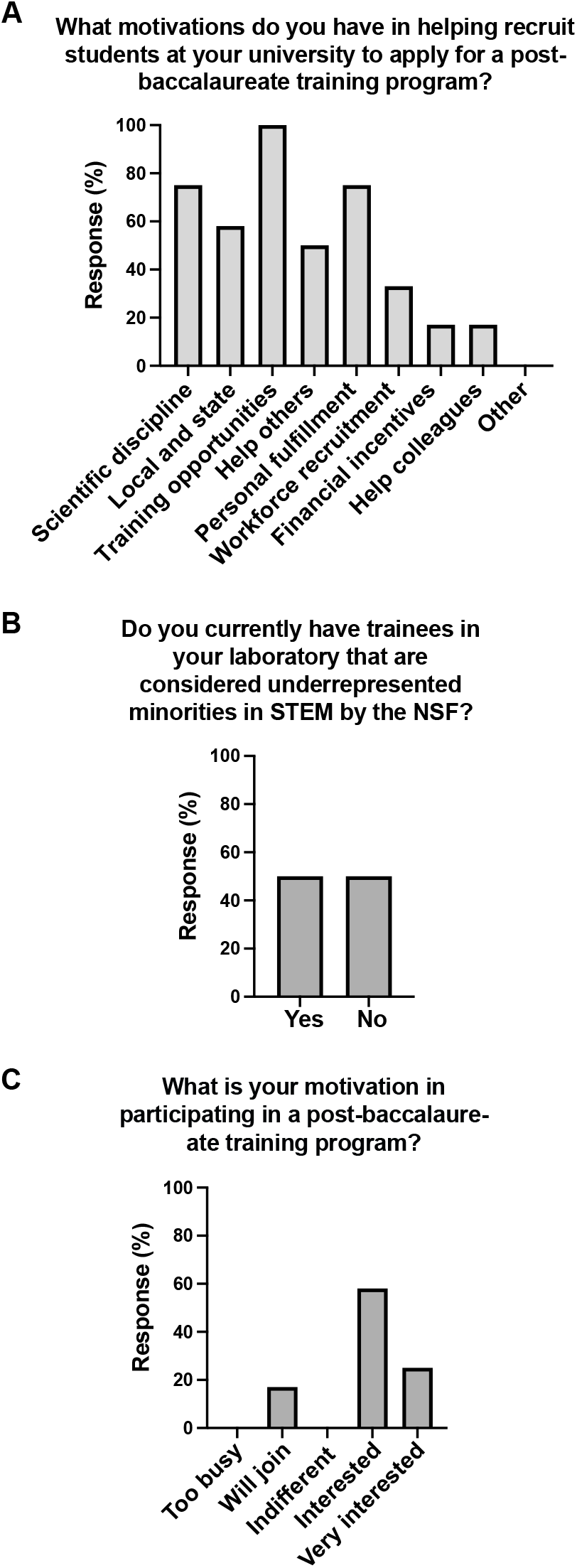
Motivation of PUI and R1 faculty to participate in a post-baccalaureate training program and their experience training underrepresented students. Post-event survey results assessing prospective mentors from participating PUI and R1 institutions for **(A)** their motivations for helping to recruit students to post-baccalaureate training programs, whether they **(B)** are currently mentoring trainees from underrepresented minority groups, and **(C)** their level of interest in participating in a post-baccalaureate training program (N = 12). **(A, C)** Response options included the following: (A) Giving back to my scientific discipline, “Scientific discipline;” giving back to my local and state community, “Local and state;” providing more training opportunities for students, “Training opportunities;” feeling of obligation to help others as I was helped, “Help others;” workforce recruitment for my lab; “Workforce recruitment;” feeling of obligation to help my colleagues, “Help colleagues.” (D) “Too busy, no interest, “Too busy;” busy, but will join if needed, “Will join;” If the opportunity arises/indifferent, “Indifferent;” very interested and actively looking for additional opportunities, “Very interested.”

**Figure 3.**
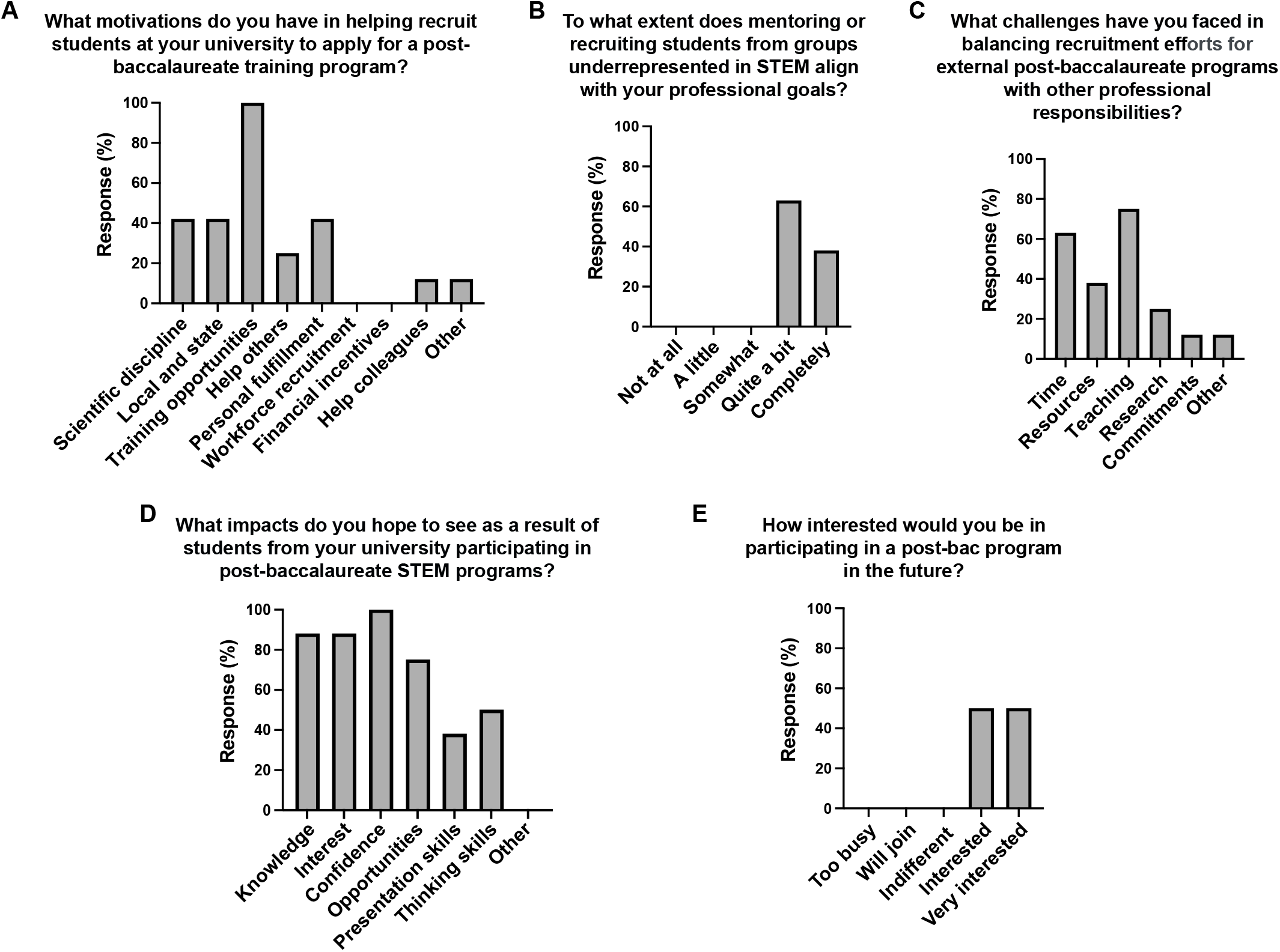
Motivations and challenges for MSI faculty in recruiting students to post-baccalaureate programs. Post-event survey results assessing STEM faculty from MSI partner institutions for **(A)** their motivations for helping to recruit students to post-baccalaureate training programs, **(B)** how well mentoring or recruiting underrepresented students in STEM aligns with their professional goals, **(C)** the challenges they face in balancing recruitment efforts with their other responsibilities, **(D)** the impacts they hope participation in post-baccalaureate programs will have on their students, and **(E)** their level of interest in participating in a post-baccalaureate training program in the future (N = 8). **(A, C, D)** Response options included the following: (A) Giving back to my scientific discipline, “Scientific discipline;” giving back to my local and state community, “Local and state;” providing more training opportunities for students, “Training opportunities;” feeling of obligation to help others as I was helped, “Help others;” workforce recruitment for my lab; “Workforce recruitment;” feeling of obligation to help my colleagues, “Help colleagues.” (C) Lack of time, “Time;” limited resources/funding, “Resources;” High teaching load, “Teaching;” research obligations, “Research;” family/personal commitments, “Commitments.” (D) Increased knowledge of research careers in STEM, “Knowledge;” increased interest in research careers in STEM, “Interest;” increased confidence in their scientific skills, “Confidence;” improved opportunities for jobs or graduate school, “Opportunities;” increased writing and speaking skills, “Presentation skills;” increased critical thinking skills, “Thinking skills.”

## Results

The goal this project was to develop an event that could recruit applicants from a range of backgrounds to post baccalaureate programs. To this end, an Open House was created to advertise a potential post-bac program to students in Indiana and the Chicago area. Partnerships were first established between three Indianapolis area PUIs and a centrally located R1 institution. Additionally, partnerships were formed with four MSIs in the Northern Indiana/Chicago area. Faculty at these MSIs interact regularly with a diverse undergraduate student body. Each MSI had a faculty contact who facilitated event advertising and chaperoned students during the travel to/from and attendance at the Open House. A full schedule of talks and social events was planned (**Fig S1**) and held at Butler University and Indiana University School of Medicine. Students learned about scientific research and professional opportunities for those entering post-bac programs and STEM careers. Discussion forums and meals were included, which allowed visiting students to discuss post-bac programs and graduate school with R1 graduate students from SACNAS and with faculty from PUI and R1 institutions.

Voluntary, anonymous pre- and post-surveys were administered at the beginning and ending of the Open House. The pre-survey solicited demographic information from the students attending the event (**Supplementary Information 1**). Information was collected regarding age, year in school, sexuality, gender, disability, military service, education, science exposure, career goals, and the attendees’ knowledge of the concept of and opportunities available in post-bac programs. All results are provided for those who responded (**Supplementary Information 1**). Of note, 14 of 15 (93%) total pre-survey respondents identified as an underrepresented racial/ethnic minority, including 7 identifying as Black/African American (47%), 7 identifying as Hispanic/Latinx/a/o/e (47%), or 1 identifying as Indigenous/American Indian or Alaskan Native (7%). Respondents were able to select multiple races/ethnicities, explaining why the number of students from minoritized groups totals more than 14. Additionally, 3 respondents (20%) indicated they had a disability according to the NIH/NSF definition (16, 17). Only 2 respondents (13%) reported having a family member in the household with a 4-year degree or higher. While all 15 respondents reported pursuit of a bachelor’s degree in science, less than half (6 respondents) could identify a science role model. A similar number (6 respondents) reported that they did not pursue independent research in their undergraduate education, either because it was not available or because they chose not to participate. The responses indicated that limited time due to work or personal obligations (9 respondents) and access to knowledge regarding research activities (7 respondents) were both significant factors in deciding whether to pursue undergraduate research. In sum, the students recruited to this Open House were members of groups typically underrepresented in science with limited exposure to science research.

Analysis of pre- and post-survey data indicated that the attending students learned about and had a positive impression of the post-bac program. Responses showed that students gained a statistically significant improvement in their understanding of post-bac training programs and what they entail after attending the Open House (**Fig 1A;** *U* (N_Pre_=15, N_Post_=13) = 26.5, *p* = 0.0003). Students also expressed a strong interest in pursuing a post-bac opportunity (**Fig 1B**). Although the pre- to post-survey gains were not statistically significant for this question [(*U* (N_Pre_=15, N_Post_=13) = 71, *p* = 0.194], this is likely due to both small sample sizes and the high number of students “agreeing” with the statement despite not being very familiar with post-bac programs in the pre-survey. Nevertheless, more students “strongly agreed” they were interested in pursuing a post-bac program in the post-survey (Mean_Pre_ = 4.07; Mean_Post_ = 4.23). The 12 post-survey responses to the question of whether the event made students more likely to apply to a post-baccalaureate program were uniformly positive. Post–event agreement ratings (median = 5, *strongly agree*) were significantly higher than a theoretical neutral median of 3 (*neither agree nor disagree*), *W* = 78, *p* = 0.0005 (two-tailed, exact; *n* = 12). All ranks were positive, indicating that participation in the Open House was consistently associated with increased agreement that students would apply to the post-baccalaureate program (**Fig. 1C**). The most positive experiences came from hearing about the benefits of a post-bac program (9 responses), an overview of a model post-bac program (9 responses), and the graduate student panel (10 responses). Anecdotally, student survey responders commented that “they definitely sold me on (the location)…and all the programs offered,” that “the event was really informative,” and that the event “was really fun and insightful. I found out more about post bac programs and the benefits.” While some students commented in the pre-survey that they were worried about the “location away from home”, “being at a predominantly white institution”, and being unsure whether completing the post-bac program “would lead to something”, none of these concerns appeared in post-survey responses. Thus, the Open House may have been successful in addressing students’ concerns. In fact, one respondent in the post-survey stated that “being away from home and finding a new place to live and having to start out my own with this change is daunting but I’m sure I’m capable of doing it.” In sum, the network-based Open House event delivered a positive experience and was successful in informing students about the benefits of a post-bac program to pursuing future careers in STEM.

PUI, R1, and MSI faculty mentors also were surveyed to assess their motivations for participating in post-bac programs. Faculty at PUIs and R1 institutions reported multiple reasons for recruiting post-bac students, including the opportunity to provide training, contribute to the scientific community, and gain personal fulfillment (**Fig 2A**). Notably, half of the survey respondents (6 of 12) are currently mentoring students from groups historically underrepresented in STEM (**Fig 2B**). Respondents also expressed sustained interest in participating in post-bac training programs (**Fig 2C**). One faculty member highlighted that some students may “realize too late that they should have done lab research in their senior year and faculty may not be willing to work with students during that short time” (**Supplementary Information 3**). Post-bac programs were seen as a valuable solution to bridge this gap. Overall, potential faculty mentors found the recruitment of post-bac students to be a rewarding experience and expressed a strong desire to remain engaged in such programs.

Survey responses from MSI faculty closely mirrored those of PUI and R1 faculty. Their primary motivation for participating in the recruiting event was to offer students additional training opportunities in STEM (**Fig 3A**), and many felt these efforts aligned with their own professional goals (**Fig 3B**), despite heavy teaching responsibilities and limited time (**Fig 3C**). Six faculty members commented that mentoring or recruiting students from groups underrepresented in STEM aligns with both their university’s mission and their personal values (**Supplementary Information 4**). Notably, one faculty member remarked they “would be just as interested in this type of program regardless of the student demographic,” indicating that some MSI participants view post-bac programs as beneficial for all students pursuing STEM careers. Faculty also expressed optimism that these programs would boost students’ confidence, interest, and knowledge about STEM pathways (**Fig. 3D**), among other benefits. Like their PUI and R1 counterparts (**Fig 2C**), MSI faculty expressed continued interest in supporting and recruiting for post-bac programs (**Fig 3E**). Overall, MSI faculty were enthusiastic about post-bac initiatives and saw them as valuable opportunities for their students’ academic and professional development.

## Discussion

Post-baccalaureate recruitment of underrepresented minorities can be challenging due to a lack of science exposure and personalized interactions. To improve outreach to underserved populations in science, an open house event was established to advertise post-bac programs to students from MSIs and surrounding universities, providing attendees first-hand interactions with faculty, staff, students, and facilities they would encounter in such programs. Students visited the campuses of a PUI and an R1 institution, heard about post-bac programs and graduate school, and had a chance to socialize with faculty and students. Pre- and post-surveys indicated that many of the students who visited represented underserved minorities in science and that the Open House both informed and left a positive impact on their impressions of post-bac programs. Faculty surveys indicated generalized support for such programs, noting their value to students’ development in STEM and a way for faculty to promote their academic mission. Hence, direct, personalized events leveraging the strengths of multiple institutions is a viable strategy to encourage trainees to pursue post-bac opportunities.

MSI partnership to enhance science outreach and development is a well-established strategy. Personal referrals are an effective means to recruit students to graduate programs (11). Furthermore, MSI partnerships have aided in recruitment of underrepresented minorities in sciences into a physical sciences graduate program (18), and encouraged participation in STEM research with the National Oceanic and Atmospheric Administration (NOAA) (19). National programs like the Leadership Alliance, comprised of 32 institutions ranging from Ivy League schools and R1s to MSIs, have been collaboratively mentoring underrepresented minority students from undergraduate through graduate training for 30 years (20). Similarly, this Open House event relied heavily on MSI faculty to recruit students through word of mouth and flyer distribution. MSI faculty members also accompanied their students to the two-day event and saw value in the program for their students. Personalized mentorship is known to enhance a student’s STEM experience and decision to enter STEM careers (21). Thus, personalized experiences, like invitations from faculty at their own institutions to an Open House event, in addition to the direct exposure to the post-bac program environment attendees receive at the Open House, are expected to increase the likelihood that students will apply to post-bac programs.

Improvements will further refine the effectiveness of the Open House. First, while MSI student participants expressed many positive sentiments regarding their experience at the event, informal conversations with student and faculty attendees indicated that they would like additional time to explore the local area, including housing options and neighborhood information, as well as a more comprehensive overview of research departments and areas, while also ensuring research talks are as accessible as possible to a wide range of students. This added time must be balanced with the limited availability of participating faculty. Second, scheduling the Open House at a time that was mutually convenient for all institutions, each with their own unique academic calendars, while also avoiding local hotel event conflicts, was challenging. Continued communication and advance planning, as well as pairing the in-person event with virtual “office hours” and other campus visits by post-bac program faculty and student representatives should minimize these challenges in the future. Third, although advertising with the partnered MSIs was effective for recruiting Open House attendees, less effort was placed on recruiting students in the area. Local students represent an additional, potentially high yield population for a post-bac program, as they would not need extensive travel to attend the Open House, and many would likely identify as an underserved minority in science. Thus, recruiting local students to post-bac programs may be extremely fruitful, as they may be comfortable committing to a program in which they know the area, universities, and faculty members involved. More effort should be made to advertise such Open House events to all students, near and far. Fourth, many students who attended the Open House event had already made career choices. Many students were interested in clinical professions, with less than half citing research as their career goal (**Supplemental Information 1**). Student mindset can change, but it may be advantageous to target college students who are undecided or leaning toward a non-clinical STEM career, as these students will be the strongest candidates for post-bac programs. Continued personalized invitations to such students from MSI, PUI, and R1 faculty, along with providing additional STEM-career focused information to candidates, will likely be most effective in achieving this goal (11). As designed, the Open House format permits flexibility for hosts to reconfigure and emphasize strengths of their geographical area, research programs, and partners to recruit their desired post-bac cohort.

## Conclusion

Overall, this work provides evidence that having in-person Open House events is an effective way to inform students, and particularly those from groups underrepresented in STEM, about post-bac programs. Post-bac programs continue to gain traction because of their strengths in preparing students for graduate school. These training opportunities are promising avenues to recruit talent from all walks of life into STEM careers. Virtual “office” hours and flyer advertising on university boards or email are affordable and can be effective for the student knowledgeable about the next steps in a STEM career. However, to recruit students unaware of the possibilities in a science career, a more active recruitment process involving experiential learning, such as an Open House event, may aid in identifying talent outside of the normal cohort and may be effective regardless of whether students have had prior research experience. This Open House model, which capitalized on the synergy of a network of partner institutions (MSIs, PUIs, and RIs), is one method for successfully identifying post-bac candidates from underrepresented groups and sharing with them the benefits of participating in a post-bac program as an integral step in their STEM career progression. Future studies could measure the degree of increased collaboration and movement of students between MSI, PUI, and R1 campuses as a result of the network model. This strategy can be modified to present the strengths of any university, training program, or geographical area. Thus, STEM training programs may consider hosting similar events to increase the diversity of their applicant cohort.

## Acknowledgments

The authors thank the students from partnering MSIs and hosting institutions for their attendance and participation. They also thank Drs. Ann Kimble-Hill, Evan Cornett, Qiuyan Chen, Emma Doud, Yangshin Park, Ms. Carmen Herrera-Sandoval, Moraima Noda, Mr. Rodney Claude, Derrick Gray, Miguel Barriera Diaz, and SACNAS (Indiana University School of Medicine (IUSM)), Center for Electron Microscopy (IUSM), and the Center for Proteome Analysis (IUSM) for speaking about their science and available post-bac programs, as well as Mr. Randall Ojeda and Ms. Mikala Lain (Butler Efroymson Diversity Center) for sharing diversity and inclusion resources and Dr. Rob Denton (Marian University) for speaking about his science. Additional thanks go out to Butler University and Indiana University School of Medicine for use of their facilities, faculty from Butler, Marian, and Indiana Universities for attending the dinner and lunch, and the Aoki Lab for help with lunch set up and clean up. Finally, the authors thank Dr. Andrew Stoehr (Butler University) for advice on statistical analysis. This project was funded by the Butler University Provost’s Office. S.T.A. is supported in part by the NIH/NIGMS (R35 GM142691).

## Figure Legends

**Supplemental Information 1**. Open House Pre-Survey Results

**Supplemental Information 2**. Open House Post-Survey Results.

**Supplemental Information 3**. PUI/RI Mentor Survey Results.

**Supplemental Information 4**. MSI Faculty Survey Results.

---

**Figure S1.**
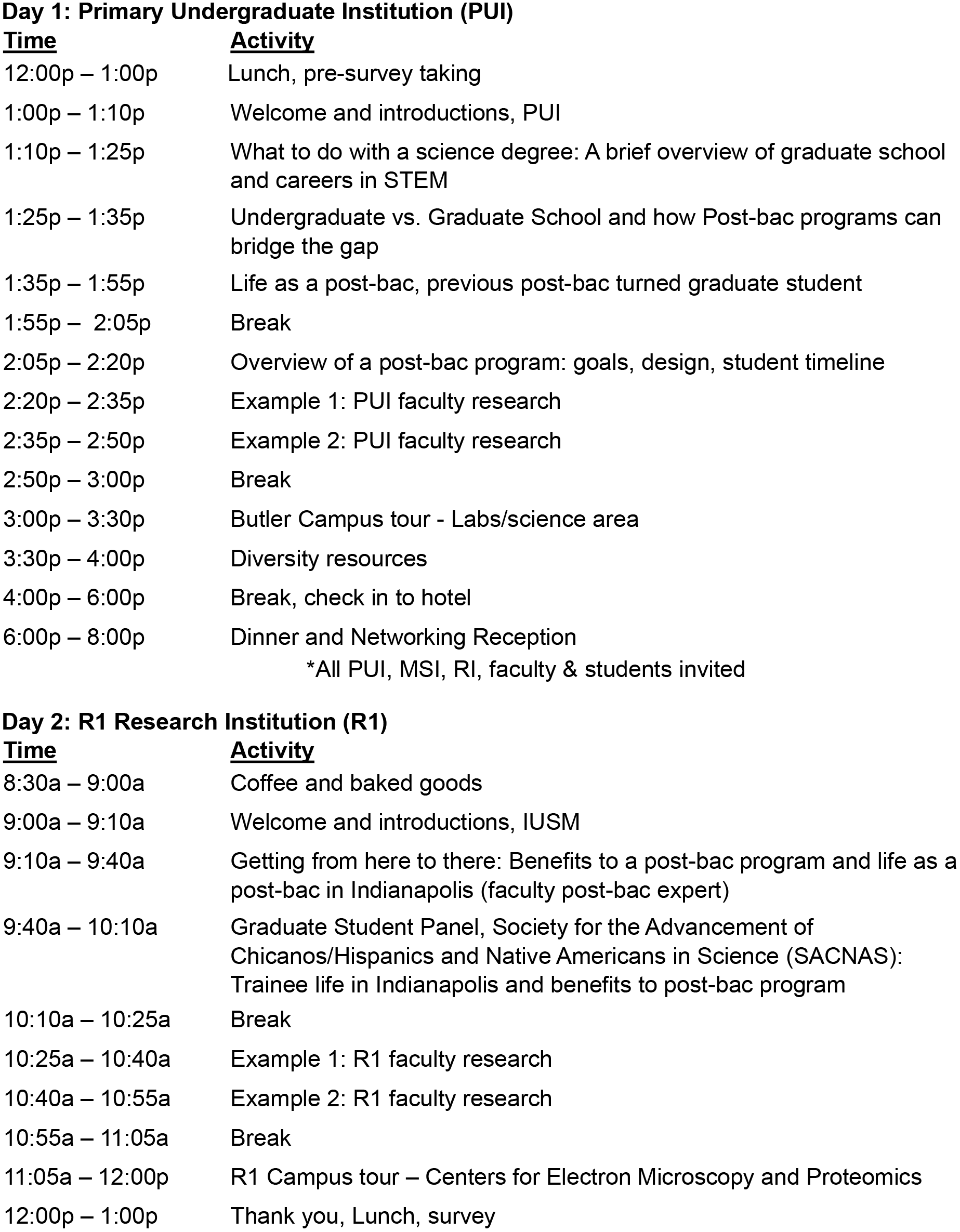
Post-Baccalaureate Open House Agenda.

## References

1. Mahatmya D, Morrison J, Jones RM, Garner PW, Davis SN, Manske J, et al. Pathways to Undergraduate Research Experiences: a Multi-Institutional Study. Innovative Higher Education. 2017;42(5):491–504. doi: 10.1007/s10755-017-9401-3.

2. Hirst RA, Bolduc G, Liotta L, Packard BW-L. Cultivating the STEM Transfer Pathway and Capacity for Research: A Partnership Between a Community College and a 4-Year College. Journal of College Science Teaching. 2014;43(4):12–7.

3. Park HY, Berkowitz O, Symes K, Dasgupta S. The art and science of selecting graduate students in the biomedical sciences: Performance in doctoral study of the foundational sciences. PLoS One. 2018;13(4):e0193901. Epub 20180403. doi: 10.1371/journal.pone.0193901. PubMed PMID: 29614110; PubMed Central PMCID: PMC5882097.

4. Postbaccalaureate Research Education Program (PREP) (R25): National Institutes of Health; 2024. Available from: https://www.nigms.nih.gov/training/PREP/Pages/default.aspx.

5. Gazley JL, Remich R, Naffziger-Hirsch ME, Keller J, Campbell PB, McGee R. Beyond Preparation: Identity, Cultural Capital, and Readiness for Graduate School in the Biomedical Sciences. J Res Sci Teach. 2014;51(8):1021–48. doi: 10.1002/tea.21164. PubMed PMID: 26366013; PubMed Central PMCID: PMC4564061.

6. Hardy TM, Hansen MJ, Bahamonde RE, Kimble-Hill AC. Insights Gained into the Use of Individual Development Plans as a Framework for Mentoring NIH Postbaccalaureate Research Education Program (PREP) Trainees. J Chem Educ. 2022;99(1):417–27. Epub 20211124. doi: 10.1021/acs.jchemed.1c00503. PubMed PMID: 36186731; PubMed Central PMCID: PMC9521764.

7. Hall A, Mann J, Bender M. Analysis of Scholar Outcomes for the NIGMS Postbaccalaureate Research Education Program 2015. Available from: https://loop.nigms.nih.gov/2015/09/outcomes-analysis-of-the-nigms-postbaccalaureate-research-education-program-prep/.

8. Schwartz NB, Risner LE, Domowicz M, Freedman VH. Comparisons and Approaches of PREP Programs at Different Stages of Maturity: Challenges, Best Practices and Benefits. Ethn Dis. 2020;30(1):55–64. Epub 20200116. doi: 10.18865/ed.30.1.55. PubMed PMID: 31969784; PubMed Central PMCID: PMC6970524.

9. Research and Mentoring for Postbaccalaureates in Biological Sciences (RaMP): National Science Foundation; 2024. Available from: https://new.nsf.gov/funding/opportunities/research-mentoring-postbaccalaureates-biological.

10. ACS Center for Diversity in Cancer Research (DICR) Training: American Cancer Society; 2024. Available from: https://www.cancer.org/research/acs-center-for-diversity-in-cancer-research-training.html.

11. Shadding CR, Whittington D, Wallace LE, Wandu WS, Wilson RK. Cost-Effective Recruitment Strategies That Attract Underrepresented Minority Undergraduates Who Persist to STEM Doctorates. SAGE Open. 2016;6(3):2158244016657143. doi: 10.1177/2158244016657143.

12. Gray M, Daugherty MK. Factors that Influence Students to Enroll in Technology Education Programs. Journal of Technology Education. 2004;15. doi: 10.21061/jte.v15i2.a.1.

13. Fischbach R. Assessing the impact of university open house activities. College Student Journal. 2006;40:227+.

14. Pew Research Center. STEM Jobs See Uneven Progress in Increasing Gender, Racial and Ethnic Diversity. 2021 April 2021. Report No.

15. Jensen-Ryan D, Murren CJ, Rutter MT, Thompson JJ. Advancing Science while Training Undergraduates: Recommendations from a Collaborative Biology Research Network. CBE Life Sci Educ. 2020;19(4):es13. doi: 10.1187/cbe.20-05-0090. PubMed PMID: 33215973; PubMed Central PMCID: PMC8693944.

16. Americans With Disabilities Act of 1990, (1990).

17. (NCSES) NCfSaES. Diversity and STEM: Women, Minorities, and Persons with Disabilities 2023. Alexandria, VA: National Science Foundation, 2023.

18. Stassun KG, Burger A, Lange SE. The Fisk-Vanderbilt Masters-to-PhD Bridge Program: A Model for Broadening Participation of Underrepresented Groups in the Physical Sciences through Effective Partnerships with Minority-Serving Institutions. Journal of Geoscience Education. 2010;58(3):135–44. doi: 10.5408/1.3559648.

19. Robinson L, Rousseau J, Mapp D, Morris V, Laster M. An Educational Partnership Program with Minority Serving Institutions: A Framework for Producing Minority Scientists in NOAA-Related Disciplines. Journal of Geoscience Education. 2007;55(6):486–92. doi: 10.5408/1089-9995-55.6.486.

20. Ghee M, Collins D, Wilson V, Pearson Jr W. The Leadership Alliance: Twenty Years of Developing a Diverse Research Workforce. Peabody Journal of Education. 2014;89(3):347–67. doi: 10.1080/0161956X.2014.913448.

21. Estrada M, Hernandez PR, Schultz PW. A Longitudinal Study of How Quality Mentorship and Research Experience Integrate Underrepresented Minorities into STEM Careers. CBE—Life Sciences Education. 2018;17(1):ar9. doi: 10.1187/cbe.17-04-0066.

